# How does V1 population activity inform perceptual certainty?

**DOI:** 10.1101/2023.09.08.556926

**Authors:** Zoe M. Boundy-Singer, Corey M. Ziemba, Olivier J. Hénaff, Robbe L. T. Goris

## Abstract

Neural population activity in sensory cortex informs our perceptual interpretation of the environment. Oftentimes, this population activity will support multiple alternative interpretations. The larger the set of plausible alternatives, the more uncertain the selected perceptual interpretation. We test the hypothesis that the reliability of perceptual interpretations can be revealed through simple transformations of sensory population activity. We recorded V1 population activity in fixating macaques while presenting oriented stimuli under different levels of nuisance variability and signal strength. We developed a decoding procedure to infer from V1 activity the most likely stimulus orientation as well as the certainty of this estimate. Our analysis shows that response magnitude, response dispersion, and variability in response gain all offer useful proxies for orientation certainty. Of these three metrics, the last one has the strongest association with the decoder’s uncertainty estimates. These results clarify that the nature of neural population activity in sensory cortex provides downstream circuits with multiple options to assess the reliability of perceptual interpretations.

## Introduction

Perceptual systems infer properties of the environment from sensory measurements that can be corrupted by nuisance variability ^1^ and neural noise ^2^. Consequently, sensory measurements are inherently ambiguous and perceptual inferences can be uncertain. This uncertainty limits the quality of perceptually-guided behavior. To mitigate this problem, uncertain observers collect additional sensory evidence ^3,4^, combine signals across sensory modalities ^5,6^, and leverage knowledge of statistical regularities in the environment ^7,8^. These behavioral phenomena all suggest that the brain can assess the uncertainty of individual perceptual inferences, as do explicit reports of confidence in perceptual decisions ^9,10^. How it does so is unknown. The optimal computational strategy (‘exact inference’) relies on knowing how stimulus and context variables drive neural activity in sensory circuits, expressing this knowledge in a generative model, and then inverting this model for a given sensory measurement ^11–13^. This operation yields a function that expresses the likelihood of every possible stimulus interpretation. The likelihood function can be used to derive the best perceptual estimate (i.e., the mode) and its uncertainty (i.e., the width). However, exact inference entails complex calculations that are intractable for many real-world perceptual tasks ^14^.

How does the brain estimate perceptual uncertainty? One appealing possibility is that it leverages certain aspects of neural activity as a direct proxy for uncertainty. For example, given that action potentials transmit information, the overall level of responsiveness of a sensory population may provide a reasonable estimate of the certainty of any inference based on that population response ^15,16^. Likewise, given that response variability and inferential uncertainty are intimately related, spatiotemporal fluctuations in neural responsiveness may provide a useful indication of perceptual uncertainty ^17–21^. And perhaps there are less intuitive aspects of sensory population activity that provide an even better indication of downstream uncertainty ^22–25^. Here, we examined these questions in macaque primary visual cortex (V1). We studied the problem of perceptual orientation estimation in the presence of nuisance variation and signal strength variability. We developed a model based decoding procedure for deriving exact inference estimates of stimulus orientation and orientation uncertainty from neural population activity. We then compared this latter estimate with different aspects of neural activity related to coding fidelity to evaluate their suitability as “candidate representations of uncertainty”.

We found that the overall strength of the population response, the cross-neural dispersion of this response, and cross-neural variability in response gain each exhibit a modest to strong association with the decoder’s orientation uncertainty. Further analysis of the relative importance of these three variables revealed that gain variability is the main driver of this association. This was true both in the presence and absence of external stimulus variability. Neural networks trained to predict orientation uncertainty from the population response reached similar performance levels as gain variability, demonstrating that our hand-picked candidate representations are effective proxies for inferential uncertainty. Together, these findings illuminate how the nature of the neural code facilitates the assessment of perceptual uncertainty by circuits downstream of sensory cortex.

## Results

### Orientation coding in V1 loses precision under high nuisance variation and low signal strength

Our approach to evaluate candidate representations of perceptual uncertainty relies on three elements. First, a rich stimulus set which elicits variable amounts of perceptual uncertainty about a stimulus feature of interest. Second, observation of the joint spiking activity of a population of sensory neurons selective for this feature. And third, a method to obtain ground truth estimates of perceptual uncertainty on a single trial basis. We reasoned that the primary visual cortex offers a fruitful test-bed for our approach. V1 neurons are selective for local image orientation ^26^ and their activity is thought to inform perceptual orientation estimates ^27–29^. V1 activity not only reflects stimulus orientation. It is also modulated by factors that impact perceptual uncertainty about stimulus orientation. These include reductions of signal strength by lowering stimulus contrast ^30,31^ and increases of nuisance variation by widening orientation dispersion ^32,33^. Building on these earlier findings, we constructed a stimulus set consisting of filtered 3-D luminance noise with filter settings varying in center direction of motion (16 levels), orientation dispersion (2 levels), and amplitude (2 levels, see Methods, Fig. 1A).

**Figure 1.**
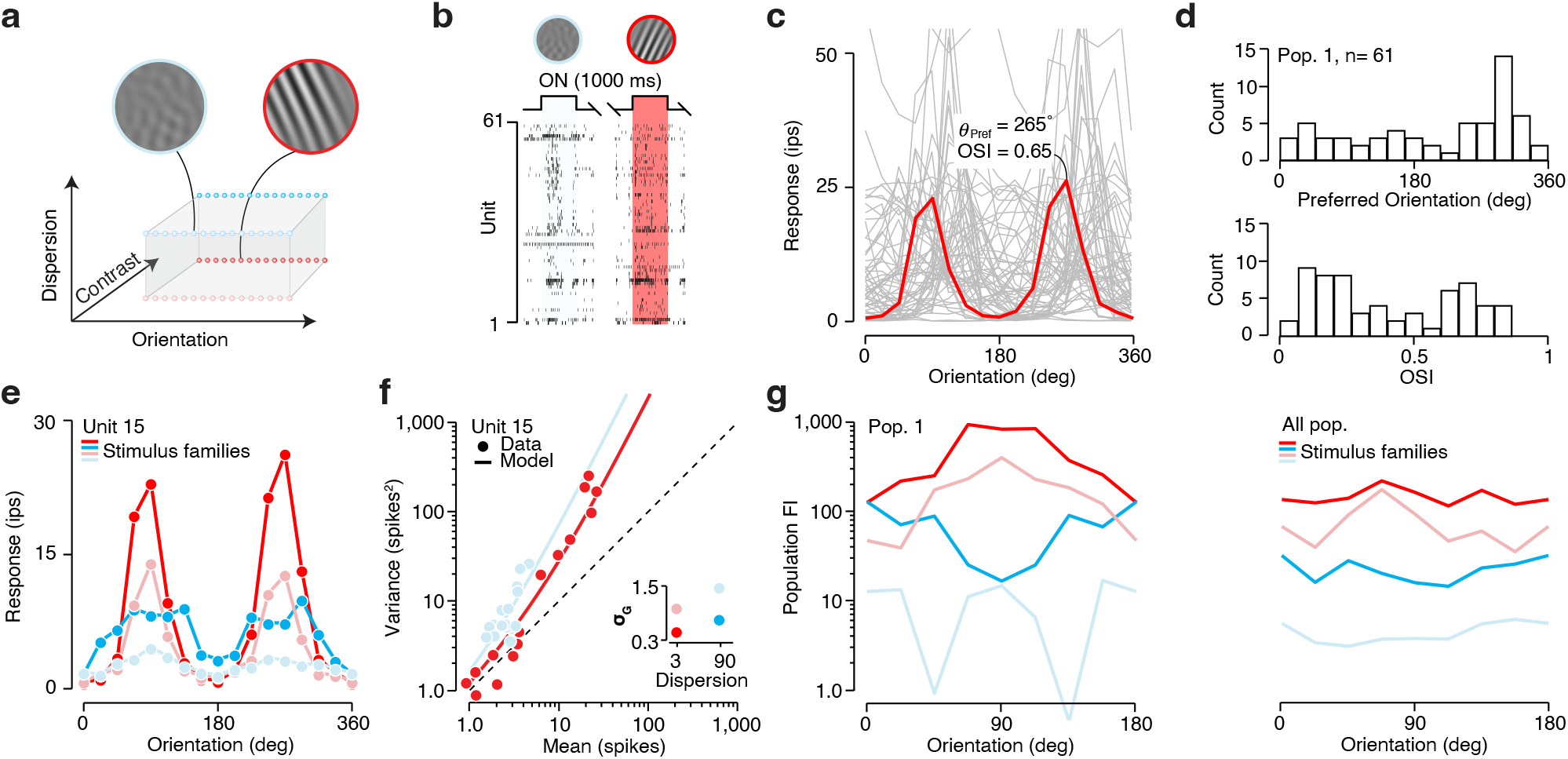
Orientation coding in V1 under different levels of nuisance variation and signal strength (**a**) We created four stimulus families, each defined by the amount of orientation dispersion and contrast energy. Each family consisted of sixteen stimuli with orientations evenly spaced from 0° to 337.5°. The example stimuli share the same orientation, but differ along the two other dimensions. (**b**) Raster plot illustrating snippets of neural activity recorded during two different trials from population 1. An example frame from each trial’s dynamic stimulus is shown on top. (**c**) Mean response to low dispersion, high contrast stimuli are plotted as a function of stimulus orientation for all units from population 1 (a single example is highlighted in red). (**d**) Distribution of preferred orientation (top) and orientation selectivity (bottom) for population 1. (**e**) Mean response as a function stimulus orientation for an example unit. Different colors indicate different stimulus families. (**f**) The example unit’s variance-to-mean relationship for two stimulus families (each data point corresponds to a different stimulus orientation). The solid line illustrates the fit of the modulated Poisson model. Inset: gain-variability, *σ*_*G*_, as estimated by the modulated Poisson model for each stimulus family for this example unit. (**g**) Population Fisher information plotted against stimulus orientation for population 1, split by stimulus family (left). Population Fisher information plotted against stimulus orientation averaged across all 13 populations (right).

We studied coding of visual uncertainty in V1 populations composed of diversely tuned units. We used linear multi-electrode arrays to record the joint spiking activity of 13 ensembles of V1 units from two fixating macaque monkeys. Ensembles ranged in size from 10 to 61 units. Stimuli were presented within a 3° wide circular window, centered on the population receptive field. We conducted these experiments in the near periphery (mean eccentricity = 2.9°), where receptive fields of V1 cells are comparatively small ^34^. Stimuli thus overlapped with both the classical receptive field and its inhibitory surround ^35^. Most units responded selectively to our stimulus set (Fig. 1B), and exhibited regular orientation tuning for the high contrast, narrowband stimuli (Fig. 1C). For each unit, we calculated the preferred direction of motion and the sharpness of its orientation tuning (see Methods). Inspection of the distribution of both statistics within each population revealed that orientation preference was typically approximately uniformly distributed (Rayleigh test for non uniformity was not significant for 10 out of 13 populations with *α* = 0.05; Fig 1D, top), and that tuning sharpness varied considerably across units (Fig. 1D, bottom).

A stimulus feature can be reliably estimated from neural population activity when that activity reliably varies with the feature. Whether or not this is the case depends on how the mean and variability of the neural response relate to the stimulus. Consider the effects of stimulus contrast and stimulus dispersion on the mean response of an example unit. Lowering contrast reduced the amplitude and, to a lesser degree, the selectivity of the response (Fig. 1E, dark vs light lines). This was true for most of our units (median amplitude reduction: 32.6%, *P* < 0.001, Wilcoxon signed rank test; median OSI reduction: 18.9%, *P* < 0.001). Increasing stimulus dispersion also reduced the example unit’s response amplitude. In addition, this manipulation substantially broadened the tuning function (Fig. 1E, red vs blue lines). Again, these effects were exhibited by most units (median amplitude reduction: 31.6%, *P* < 0.001; median OSI reduction: 64.2%, *P* < 0.001).

Our stimulus manipulations also impacted cross-trial response variability. Neurons in visual cortex typically exhibit super-Poisson variability, meaning that spike count variance exceeds spike count mean (Fig. 1F, symbols). This behavior is well captured by a statistical model of neural activity in which spikes arise from a Poisson process and response gain fluctuates from trial to trial ^36,37^ (the modulated Poisson model, Fig. 1F, lines). The larger the gain variability, *σ*_*G*_, the larger the excess spike count variance. For each unit, we fit the modulated Poisson model separately per stimulus family (i.e., a specific combination of stimulus contrast and orientation dispersion). Lowering contrast and increasing dispersion both increased gain variability (median increase: 27.1%, *P* < 0.001 for contrast, and 12.8%, *P* < 0.001 for dispersion, Wilcoxon signed rank test), consistent with previous observations in anesthetized animals ^20^.

To quantify the collective impact of these effects on the population representation of stimulus orientation, we computed the population Fisher Information (*I*_*θ*_, see Methods). This statistic expresses how well stimulus orientation can be estimated from population activity by an optimal decoder and is inversely related to the uncertainty of this estimate ^38^. Consider the Fisher Information profiles of an example population. Reducing stimulus contrast and increasing orientation dispersion both lowered *I*_*θ*_ in a systematic manner, as is evident from the vertical separation of the colored lines (Fig. 1G, left). These effects were present in all of our recordings (Fig. 1G, right), though the exact impact differed somewhat across populations. Trial-to-trial response fluctuations are often correlated among neurons ^39^. These so-called ‘noise correlations’ can affect the coding capacity of neural populations ^40,41^. However, we found no systematic effect of our stimulus manipulations on the average strength of pairwise response dependencies (*P* = 0.61 for contrast, Wilcoxon rank sum test and *P* = 0.10 for dispersion; Fig. S1A). Moreover, a shuffling analysis revealed that noise correlations had minimal impact on Fisher Information estimates (see Methods; Fig S1B-C). This result may in part be due to the relatively small sizes of our populations ^40^. Nevertheless, we conclude that under these experimental conditions, our stimulus reliability manipulations substantially impact orientation coding in V1 because they alter the relation between stimulus orientation on the one hand, and response mean and response variance on the other hand.

### The width of the likelihood function captures coding fidelity at the single trial level

To evaluate candidate representations of uncertainty, we need to know which trials yielded a high quality representation of stimulus orientation and which did not. Population Fisher information cannot be used for this because its calculation takes into account response variation across trials. Instead, we developed a decoding method to infer stimulus features from population activity on a trial-by-trial basis ^15,25,42–46^. Specifically, we built a stimulus-response model that describes how our stimulus set drives spiking activity of V1 neurons and then inverted this model to obtain an estimate of the likelihood of every possible stimulus orientation on the basis of a single trial population response. We described each unit’s responses with a previously proposed model of V1 function (the stochastic normalization model ^20,47^, see Methods; Fig. S2). In this model, stimuli are processed in a narrowly tuned excitatory channel and a broadly tuned inhibitory channel and spikes arise from a modulated Poisson process ^20,32^. As can be seen for an example unit, the model captures how both spike count mean and spike count variance depend on stimulus orientation, stimulus contrast, and stimulus dispersion (Fig. 2A). To evaluate the model’s goodness-of-fit, we calculated the association between predicted and observed response mean and variance (Fig. 2B). The association was generally high (median Pearson correlation across all units was 0.83 for response mean and 0.80 for response variance; Fig. 2C). For each unit, the model predicts how probable a given response is for a given stimulus condition. Conversely, knowledge of this probabilistic relation allows inferring how likely a given stimulus is in light of an observed response ^48^. Computing this value for all possible stimuli yields a likelihood function (see Methods). Intuitively, this function summarizes which stimulus interpretations can be ruled out and which cannot. The likelihood function provides a framework to study neural coding that naturally generalizes to the population level. If neural responses are statistically independent, the joint likelihood function of a population of neurons is simply given by the product of the likelihood functions of all units. Given that noise correlations were generally small and did not significantly impact orientation coding capacity in our data (Fig. S1), we opted to use this formulation ^20^ (see Methods).

**Figure 2.**
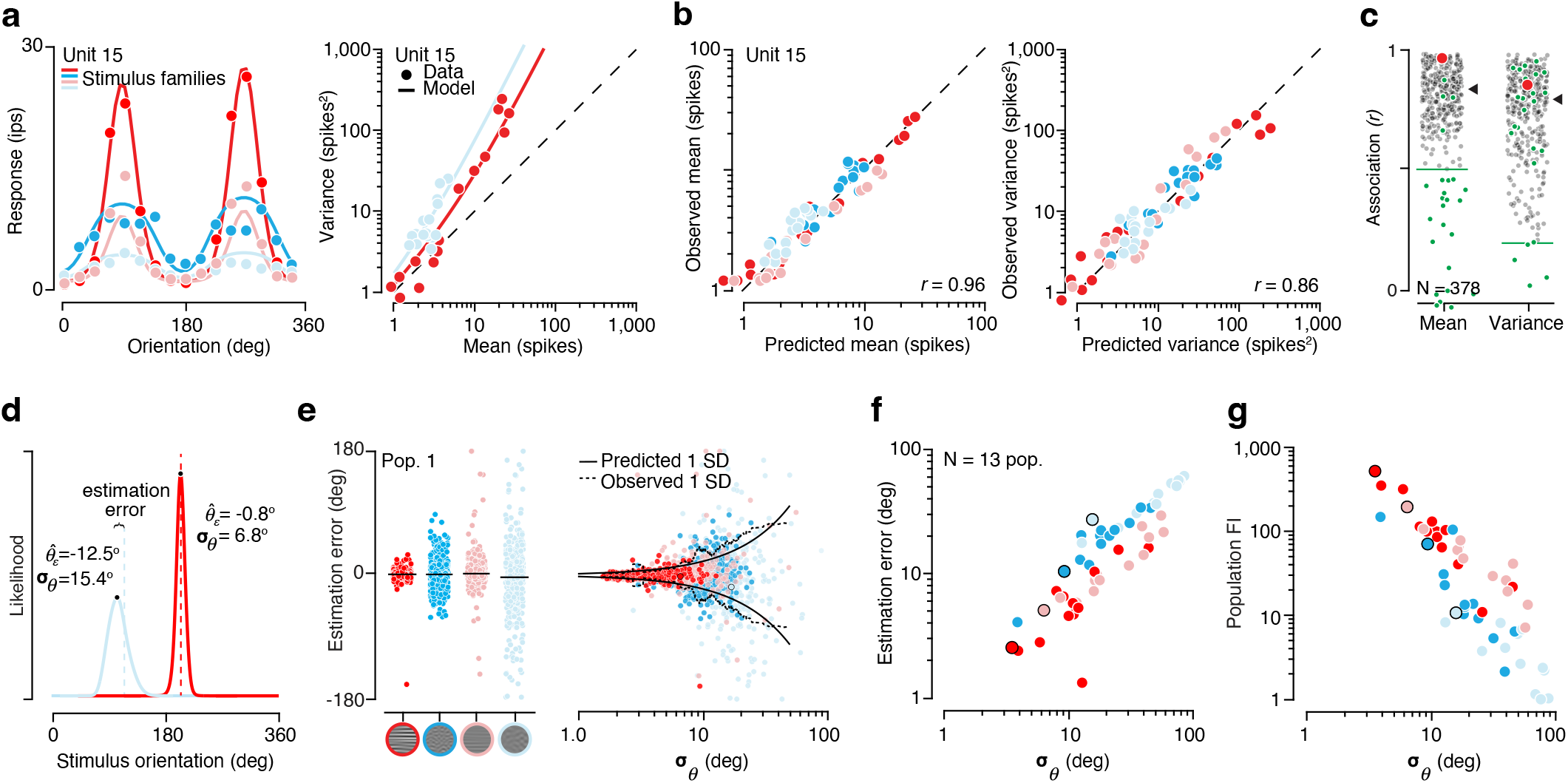
Quantifying orientation uncertainty at the single trial level (**a**) Left, mean response of an example unit as a function of orientation for all four stimulus families. Right, response variance plotted as a function of response mean for two stimulus families (one data point per orientation). Solid lines indicate the fit of the stochastic normalization model. (**b**) Observed vs predicted response mean (left) and response variance (right) for the example unit. (**c**) Distribution of the association between predicted and observed response mean (left) and response variance (right) across all units. Black triangles indicates median value across all units. Green line indicates the inclusion criterion. Units were included if the association between predicted and observed mean and variance exceeded both criteria. Excluded units are indicated with green circles. The red circle indicates the example unit illustrated in panel a,b. (**d**) Likelihood functions (solid lines) for two example trials (blue, low-contrast high-dispersion stimulus; red, high-contrast low-dispersion stimulus). Dashed line indicates the stimulus orientation. The black circle indicates the maximum likelihood orientation estimate. (**e**) Left, estimation error split by stimulus family for one example population. Each dot indicates a single trial. Black lines indicate the median estimation error. Right, estimation error as a function of likelihood width. Each dot indicates a single trial; color indicates stimulus family. Solid black line indicates the theoretically expected relationship between the variance of estimation error and likelihood width. Dashed line indicates empirically observed variance of estimation error calculated as a moving standard deviation of 100 trials sorted by likelihood width. (**f**) Average absolute estimation error per stimulus family and population as a function of the average likelihood width. Observations from population 1 are highlighted with a black outline. (**g**) Population Fisher information plotted against average likelihood width, same conventions as panel f.

Consider the likelihood function for two example trials. This function was obtained by marginalizing the full likelihood function over the dimensions of stimulus contrast and stimulus dispersion (see Methods). Population activity differed substantially across these two trials (Fig. 1B). Accordingly, the corresponding likelihood functions also differed in central tendency and shape (Fig. 2D). For each trial, we used the maximum of the marginalized likelihood function as a point estimate of stimulus orientation. Overall, this estimate correlated well with the veridical stimulus orientation (circular correlation computed across all stimulus conditions, r = 0.76 on average and ranged from 0.96 for high contrast low dispersion stimuli to 0.43 for low contrast high dispersion stimuli, Fig. S3A-B). However, it did not track stimulus orientation perfectly. On some trials, the orientation estimation error could be substantial, especially when stimulus dispersion was high (Fig. 2E, left). This pattern reflected an underlying structure in the distribution of estimation error. Specifically, as expected from a well-calibrated model, the spread of the estimation error approximately matched the width of the likelihood function (Fig. 2E, right). Consequently, the average width of the likelihood function was strongly associated with the average size of the estimation error (Spearman’s rank correlation coefficient: r = 0.91, p < 0.001; Fig. 2F) and with population Fisher Information (Spearman’s rank correlation coefficient: r = -0.81, p < 0.001; Fig. 2G). Together, these results establish the width of the likelihood function calculated under the stochastic normalization model as a principled metric of coding quality for our data set. In the following analyses, we will use it as the “ground truth” estimate of stimulus uncertainty on a trial-by-trial basis.

### Some aspects of population activity predict the decoder’s stimulus uncertainty

Might certain transformations of neural activity in sensory cortex provide a direct proxy for perceptual uncertainty? Motivated by previous theoretical proposals ^15,16,19,20,49^, we evaluate three different aspects of V1 population activity as candidate representations of coding quality: (1) the overall strength of the population response (‘response magnitude’), (2) cross-neural variability in responsiveness (‘response dispersion’ as absolute measure, and ‘relative dispersion’ as normalized measure), and (3) variability in excitability (‘gain variability’). For each metric, we consider two variants. One that can be estimated directly from the observed population response (‘direct estimates’), and one that takes into account knowledge of the units’ stimulus-response relation (‘inferred estimates’, Fig. 3A). We selected these metrics because they each capture an aspect of neural activity that is thought to either influence or reflect coding fidelity. Specifically, given that cortical responses appear subject to Poisson-like noise ^30,36,50^, response magnitude is a proxy for the signal-to-noise ratio of the neural response. Cross-neural response variability, on the other hand, in part reflects the selectivity of the stimulus drive. It will only be high for trials in which a subset of neurons is highly active – an activation pattern that allows for unambiguous stimulus inference ^48^ and that satisfies the goal of coding efficiency ^49^. Finally, it has been shown previously that cross-trial gain variability in V1 tracks orientation uncertainty within single units ^20^ (also see Fig. 1F, inset). This relationship may generalize to cross-neural gain variability within populations.

**Figure 3.**
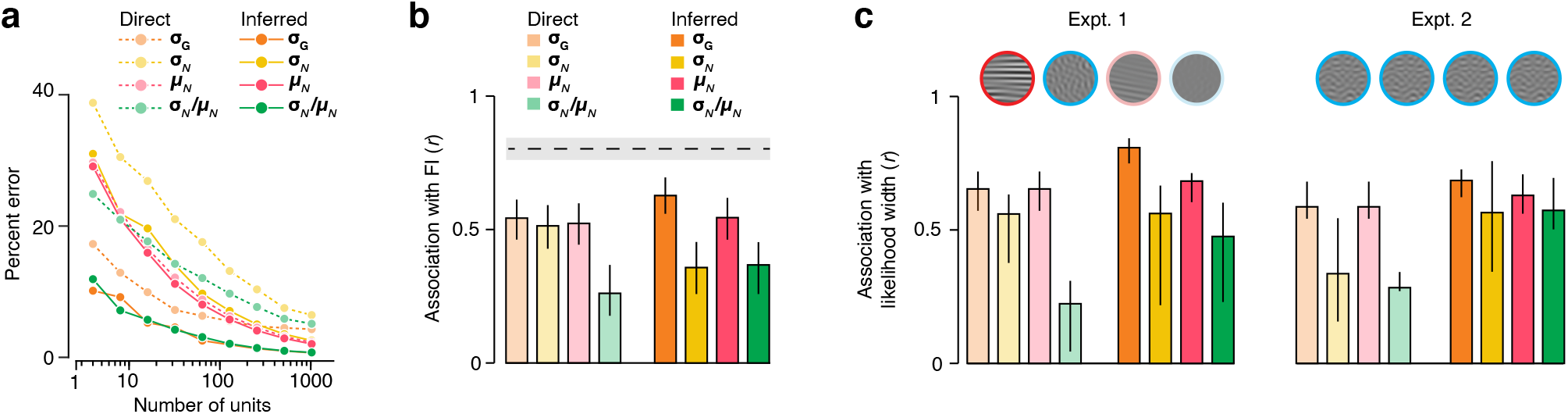
Evaluating different candidate metrics of uncertainty (**a**) Percent error in recovery of the ground truth value of each direct and inferred metric for simulated populations of various sizes. (**b**) Spearman rank correlation between candidate uncertainty metrics and population Fisher information. Light colors correspond to direct metrics; dark colors correspond to inferred metrics. Error bars indicate 75% confidence intervals of bootstrapped correlation values. The dashed line indicates the rank correlation between likelihood width and Fisher information with 75% confidence interval of bootstrapped correlation values indicated with a gray band. (**c**) Left/top, example sequence of stimuli for experiment 1. Stimuli randomly varied in orientation, family, and noise seed. Left/bottom, bar plot indicating the association between candidate neural metrics of uncertainty and likelihood width. Right/top Example sequence of stimuli for experiment 2. A single orientation, family, and noise seed defined a stimulus that was over-represented relative to all other stimulus conditions. Only trials in which this stimulus was shown were analyzed. Right/ bottom, bar plot indicating the association between candidate neural metrics of uncertainty and likelihood width for these trials. Error bars indicate interquartile range.

We reasoned that proxies for perceptual uncertainty ought to meet three requirements to be useful: (1) distinguish reliable from unreliable stimulus conditions; (2) predict perceptual uncertainty on a trial-by-trial basis; and (3) predict variability in perceptual uncertainty solely due to internal sources. Motivated by this logic, we first examined the relationship between the selected metrics and Fisher Information. To this end, we computed the average value of each metric for each stimulus family, just like we did previously for the width of the likelihood function (Fig. 2F,G). Consider the direct estimates. Response magnitude, response dispersion, and gain variability each exhibited a modest association with Fisher information (r = 0.52, 0.51, and 0.54, with 95% confidence intervals ranging from 0.31-0.69, 0.30-0.69, and 0.35-0.70; Fig. 3B, left), but fell short of the predictive power of the width of the likelihood function (r = 0.80, 95% confidence interval 0.68-0.88). Note that we consider two measurements statistically different at a P-value of 0.05 when the observed value for one does not fall within the other’s 95% confidence interval. By comparison, relative dispersion had a substantially weaker correlated with Fisher information (r = 0.26, 95% confidence interval 0.02-0.47; Fig. 3B, left). This pattern may in part reflect the limited size of our recorded populations. For each of these metrics, simulated direct estimates better approximate the ground truth as population size grows (Fig 3A). However, the speed of improvement differs across metrics, implying that some will be more hampered by our recording conditions than others (Fig 3A). We therefore complemented this analysis with one in which we attempted to get better estimates of each metric by inferring them from the observed population activity whilst taking into account knowledge of each unit’s tuning properties (see Methods). In simulation, this procedure improves estimation accuracy for each metric (Fig 3A, full vs dotted lines). Inferred response magnitude and gain variability each exhibited a modest association with Fisher information (r = 0.55 and 0.63, with 95% confidence intervals 0.35-0.71 and 0.45-0.77) while response dispersion and relative dispersion tended to exhibited a weaker association (r = 0.36 and 0.37, with 95% confidence intervals 0.12-0.57 and 0.13-0.56).

We next asked how well these metrics predict perceptual uncertainty on a trial-by-trial basis. For each population, we quantified this association by computing the rank correlation between the metric under consideration and the width of the likelihood function. We summarized results across populations by computing the mean rank correlation. This yielded values that were modest to high for inferred response magnitude (r = 0.68, 95% confidence interval of the mean 0.54-0.83), response dispersion (r = 0.54, 0.36-0.72), and gain variability (r = 0.81, 0.74-0.84), but not for relative dispersion (r = 0.46, 0.27-0.64; Fig. 3C, left).

Perceptual uncertainty in part arises from internal sources ^2,14^. To isolate variations in this factor, we conducted a variant of our experiment in eight recordings in which a single, randomly chosen stimulus was over-represented and shown several hundred times over the course of the experiment. As expected, this set of identical trials elicited considerable variability in the width of the likelihood function (standard deviation = 19.75 deg on average, 38.9 % smaller than the standard deviation across all other trials in these experiments; see Methods). This variability must be largely due to internal sources. We found that it was predicted similarly well by all four inferred candidate metrics (mean rank correlation r = 0.63, 95% confidence interval 0.49-0.77 for response magnitude, 0.54, 0.30-0.77 for response dispersion, 0.69, 0.58-0.79 for gain variability, and 0.55, 0.25-0.82 for relative dispersion; Fig. 3C, right).

In summary, our analysis suggests that response magnitude, response dispersion, and gain variability all provide a useful proxy for the fidelity of orientation coding in V1. This was true both in the presence and absence of external sources of stimulus variability. By the same standard, the combination of two candidate metrics in the form of relative dispersion did not result in a more predictive metric and thus not does not appear to be a useful proxy for stimulus uncertainty. These results do not appear to depend on the details of the decoding method. Specifically, decoding V1 activity with a conceptually simpler method that treats spikes as if they arise from a pure Poisson process yielded qualitatively similar results, though the association between the candidate metrics and likelihood width were overall weaker (Fig. S4).

### Gain variability directly predicts stimulus uncertainty, the other metrics do not

We have identified three candidate metrics that exhibit modest to strong correlations with the width of the likelihood function. This implies that they may also correlate strongly with each other. Indeed, the average rank correlation between inferred response magnitude and response dispersion was 0.91. For response magnitude and gain variability it was 0.86 and for response dispersion and gain variability it was 0.68. We wondered how controlling for this confound would impact the metrics’ association with stimulus uncertainty. To this end, we rank ordered all trials from a given recording as a function of the metric we sought to control for. We then selected the first 50 trials. By design, these trials will not vary much along the sorting metric (i.e., the ‘frozen’ variable), but they may vary along the two other ‘non-frozen’ metrics (Fig. 4A, left). The correlation between the non-frozen metrics and the width of the likelihood function provides a measure of the strength of the association in the absence of a confounding variable (Fig. 4A, right). We then repeated this analysis for the next sets of 50 trials and thus obtained a distribution of this measure (see Methods). Controlling for response magnitude had a modest impact on the predictive value of gain variability (mean rank correlation: r = 0.53, for experiment 1, and 0.32, for experiment 2, Fig. 4B,C). Conversely, freezing gain variability all but nullified the association between response magnitude and likelihood width. This was true of experiment 1 (r = 0.04, difference with gain variability: P < 0.001, Wilcoxon Rank Sum Test; Fig. 4B, left) and experiment 2 (r = 0.11, difference with gain variability: P < 0.001; Fig. 4C, left). We found a similar asymmetric pattern for gain variability and response dispersion. Freezing response dispersion did little to the predictive power of gain variability (r = 0.63 for Experiment 1, and 0.40 for experiment 2, Fig. 4B,C), but freezing gain variability removed most of the association between response dispersion and likelihood width (r = 0.04 for Experiment 1, difference with gain variability: P < 0.001; r = 0.10 for experiment 2, P < 0.001; Fig. 4B,C middle). This data pattern suggests that out of these three candidate metrics, gain-variability has the most direct association with stimulus uncertainty. A complementary analysis that sought to examine how each candidate metric’s correlation with likelihood width depended on the inter-metric correlation further corroborated this conclusion (see Fig. S5).

**Figure 4.**
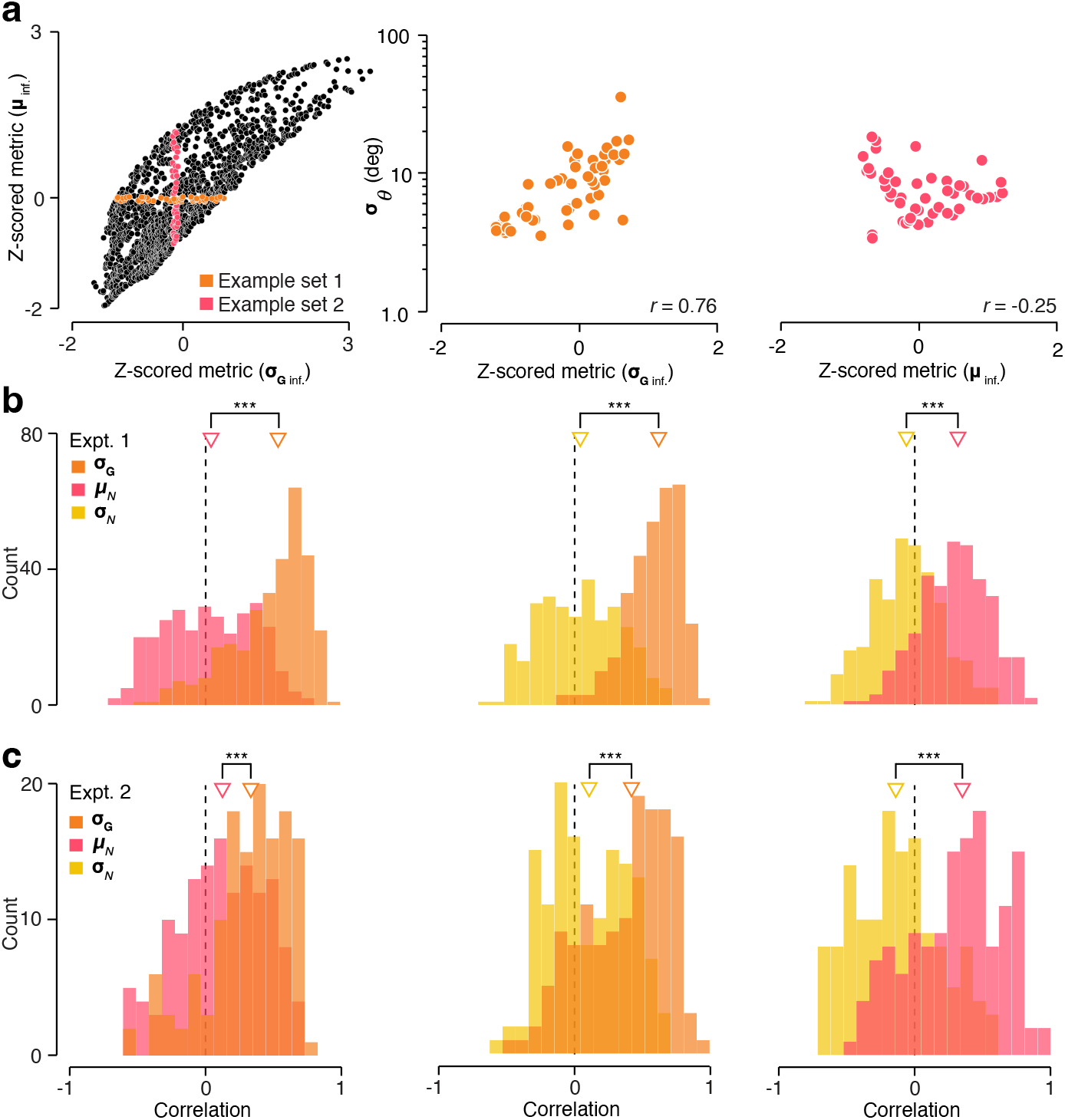
Comparison of partial correlations between three candidate metrics and the decoder’s stimulus uncertainty. (**a**) Left, population rate is plotted against gain-variability for an example population. Each point represents a single trial. Colored points indicate 50 consecutive rank-sorted trials. The two highlighted example sets of data points have identical variance in the non-frozen metric (*σ*^2^ = 0.35). Middle, relationship between gain variability and likelihood width for an example set of trials in which the population rate was kept constant (i.e., the orange trials in the left panel). Right, relationship between population rate and likelihood width for an example set of trials in which gain variability was kept constant (i.e., the pink trials in the left panel). (**b**) Distribution of partial correlations across all experiment 1 recordings for population rate and gain variability (left), response dispersion and gain variability (middle), and response dispersion and population rate (right). (**c**) Same as panel b for Expt 2. Triangles represent mean correlation; *** P < 0.001, Wilcoxon Rank Sum Test.

### Gain variability rivals performance of artificial neural networks

Our choice of candidate metrics was motivated by previous theoretical proposals ^15–17,19,20,49^. It is of course possible that there exists a hitherto unsuspected transformation of population activity that provides an even better indication of the decoder’s uncertainty. To complement our analysis with a more agnostic approach, we trained a set of artificial neural networks (ANNs) to predict the width of the likelihood function from the population response. ANNs differed in architecture and training regime. To identify the subset of ANNs most relevant to our purposes, we conducted a two-round analysis. In the first round, we considered a diverse set of ANNs which differed in their architecture (specifically, the number of hidden layers and hidden units) and training regime (specifically, the dropout rate and weight decay, see Methods). For each population, we selected the combination of network architecture and training regime that yielded maximal predictive accuracy on held out data (see Methods). We found that the best-performing combinations differed across the 13 populations. In the second round, we therefore again considered a set of ANNs, but this time the variation in network architecture and training regime was limited to the cases that had at least once resulted in a best performance in round 1. We report the average performance across networks as figure of merit. As we did previously for the hand-picked metrics, we first computed the correlation between the average predicted uncertainty for each stimulus family and population Fisher Information. This association rivaled the best handpicked metric, inferred gain variability (r = 0.61 vs 0.63, with 95% confidence intervals ranging from 0.59-0.62 and 0.45-0.77). We next examined how well the ANNs predicted likelihood width on a trial-by-trial basis. For experiment 1, we found that likelihood width was on average not significantly better predicted by the family of ANNs than by inferred gain variability (mean r = 0.79 with 95% confidence intervals 0.75-0.83 for ANN predicted likelihood width, mean r = 0.81, 0.69-0.90 for inferred gain variability; Fig. 5B, left). For experiment 2, we found that likelihood width was somewhat better predicted by ANNs than by inferred gain variability (mean r = 0.77 with 95% confidence intervals 0.71-0.83 for ANN predicted likelihood width, mean r = 0.69, 0.58-0.81 for inferred gain variability; Fig. 5B, right). We conclude that gain variability captures much of the variance of perceptual uncertainty that can be captured by a simple transformation of sensory population activity.

**Figure 5.**
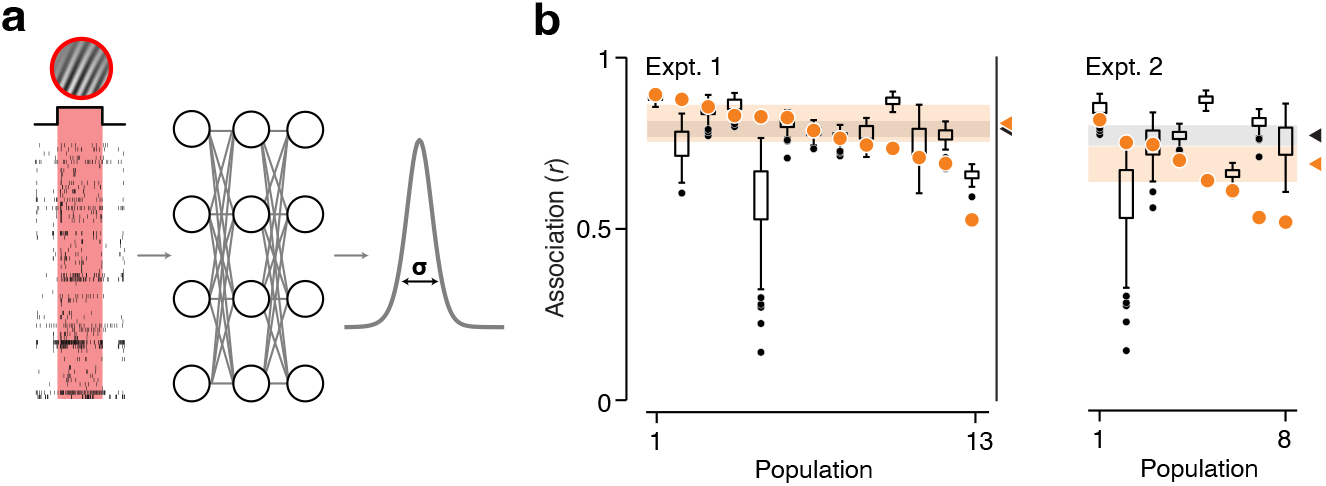
Artificial neural network prediction of likelihood width. (**a**) ANNs were trained on per-trial vectors of spike counts to predict the width of the likelihood function. (**b**) Boxplots indicate association of network predicted and ground truth likelihood width per population across the set of networks for experiment 1 (left) and experiment 2 (right). Box indicates interquartile range. Solid black lines indicate the central 95% of the distribution. Black dots indicate outlier networks. Orange dots indicate inferred gain variability association with likelihood width. Gray and orange shaded regions indicate the 68% confidence interval around the mean association between network or metric and likelihood width. Triangles indicate mean association between network or metric and likelihood width.

## Discussion

Uncertainty is intrinsic to perception. Inevitably, some perceptual interpretations of the environment are more uncertain than others. A prominent hypothesis holds that the same sensory neurons that inform perceptual interpretations represent the uncertainty of these interpretations ^15–22,25,45^. The nature of this representation is debated. A major challenge in studying this is that perceptual uncertainty is a property of an observer’s belief about the world, not a property of the world itself ^51^. Here, we sought to overcome this by manipulating stimulus reliability in two distinct ways and by using a model-based procedure to decode V1 spiking activity as a stand-in for what downstream areas *ought to* believe about stimulus orientation. We found that response magnitude, response dispersion, and variability in response gain all offer useful proxies for the certainty of stimulus orientation estimates. This was also true when fluctuations in uncertainty were not due to external factors but instead arose from internal sources. Of the metrics we considered, gain variability exhibited the most direct association with stimulus uncertainty.

Our findings offer empirical support for the central tenets of two different theoretical frameworks for the representation of uncertainty in sensory cortex: probabilistic population codes (PPC) and the sampling hypothesis. The PPC framework is built on the idea that the nature of the neural code should enable implementing statistical inference with simple mathematical operations in a feedforward manner ^16,22,52^. The modest to strong association of three simple transformations of population activity with an optimal decoder’s uncertainty estimates validates this notion for the primary visual cortex. The core idea of sampling models, on the other hand, is that an aspect of response variability is used to represent uncertainty ^17–19,21^. The finding that gain variability is the purest predictor of stimulus uncertainty further corroborates this hypothesis for the primary visual cortex ^19–21^. Interestingly, recent theoretical work has shown that some coding regimes are consistent with both the PPC and sampling framework ^13^. It is possible that the primary visual cortex operates in exactly this mode ^13^.

From a statistical standpoint, the three metrics we deemed useful proxies for stimulus uncertainty capture very different aspects of neural population activity: the average of the neural response, the dispersion of these responses, and the variability of a latent modulator in a doubly stochastic process. Empirically, we found these metrics to be highly correlated with each other. Why might this be so? We think that response average and response dispersion are closely related because spike counts in visual cortex tend to be exponentially distributed ^53^. In principle, this distribution allows maximal information transmission per spike ^54–56^. Under the exponential distribution, response standard deviation is equal to the response mean, providing an explanation for the strong association between both statistics in our data. We further suggest that response average and gain variability are closely related because of the mechanistic origins of gain variability. Specifically, we speculate that gain variability in large part arises from stimulus-independent noise in a divisive normalization signal. This normalization signal is thought to reflect aggregated nearby neural activity ^47,57^. As shown previously, noisy normalization naturally results in gain variability being inversely proportional to the normalization signal ^20,58^. It follows that a higher mean response, leading to a stronger normalization signal, will generally coincide with a lower level of gain variability. If these interpretations are correct, then these strong associations might not just be specific to this V1 experiment but reflect a general property of visual cortex.

Regardless of its source, the strong association between the different statistics we studied implies that downstream circuits have multiple options to assess the reliability of the sensory messages conveyed by neural populations in visual cortex. Perhaps a simple transformation is favored in cases where an approximate uncertainty estimate suffices, and more complex transformations are used when achieving a goal critically depends on the quality of the perceptual certainty estimate. More generally, we expect that the quality of perceptual certainty estimates will improve with experience, as is the case for simple perceptual decisions ^59^. Investigating these questions requires measuring sensory population activity from an animal generating a behavior that directly reflects their perceptual certainty on a trial-by-trial basis ^25,60,61^.

The metrics we considered as candidate representations of uncertainty can be estimated directly from neural population activity without knowledge of the tuning properties of the neurons or of the selected stimulus interpretation. However, for these estimates to be reliable, relatively large populations are required. We therefore resorted to an estimation technique that leveraged knowledge of stimulus response relations. While this approach is principled, verifying whether these results hold for direct estimation when the recorded population size is substantially larger is an important goal for future work.

## METHODS

### Physiology

All electrophysiological recordings were made from two awake fixating adult male rhesus macaque monkeys (*Maccaca mulatta*, both 7 years old at the time of recording). Subjects were implanted with a titanium chamber ^62^ which enabled access to V1. All procedures were approved by the University of Texas Institutional Animal Care and Use Committee and conformed to National Institutes of Health standards. Extracellular recordings from neurons were made with one or two 32-channel S probes (Plexon), advanced mechanically into the brain with Thomas recording microdrives. Spikes were sorted with the offline spike sorting algorithm Kilosort2^63^, followed by manual curation with the ‘phy’ user interface (https://github.com/kwikteam/phy). An example snippet of neural activity is shown in Fig. 1B.

### Apparatus

Headfixed subjects viewed visual stimuli presented on a gamma-corrected 22-inch CRT monitor (Sony Trinitron, model GDM-FW900) placed at a distance of 60 cm. The monitor had a spatial resolution of 1280 by 1024 pixels and a refresh rate of 75 Hz. Stimuli were presented using PLDAPS software ^64^ (https://github.com/HukLab/PLDAPS).

### Stimuli

Stimuli consisted of band-pass filtered 3-D luminance noise. The filter was organized around a tilted plane in the frequency domain which specified a particular direction and speed of image motion (i.e., it was velocity-separable). All stimuli had a central spatial frequency of 2.5 cycles/deg with a bandwidth of 0.5 octaves and a central temporal frequency of 2.5 deg/sec with a bandwidth of 1 octave. Orientation bandwidth was either 3° or 90°, corresponding to the “low” and “high” level of dispersion. Stimulus contrast was computed by normalizing the summed orientation amplitude spectrum of each stimulus frame with the summed amplitude spectrum of a reference grating with matching spatial frequency. We equated contrast across orientation bandwidths by rescaling the filter output appropriately. The stimulus set was composed of four stimulus families (two orientation dispersion levels × two contrast levels), which each contained 16 differently oriented stimuli, evenly spaced between 0° and 337.5°. For each stimulus condition, we generated five unique stimulus versions by using a different noise seed. Stimuli were presented within a vignette with a diameter of 3°, centered on the estimated average receptive field location, determined through a hand-mapping procedure. Stimuli were presented in random order for 1,000 ms each. Trials were excluded from further analysis if fixation was not maintained within a radius of 0.8 degrees from the fixation point for the duration of the stimulus presentation. The number of repeats per condition varied from session to session. In experiment 1 the average number of repeats per stimulus condition was 28.3 ± 5.6. In experiment 2 one stimulus was randomly selected to be over-represented. In these recordings the average number of repeats for non-selected stimuli and the single randomly selected stimulus was 16.9 ± 2.1 and 1089 ± 98.9 respectively.

### Data Analysis: single units and unit pairs

We first studied the units’ orientation tuning in response to the high-contrast low-dispersion stimuli (examples shown in Fig. 1C). For each unit, we chose a response latency by maximizing the stimulus-associated response variance ^65^, and counted spikes within a 1,000 ms window following response onset. We visually inspected tuning curves and excluded untuned units from further analysis (27 % of units). For each unit, we estimated the preferred stimulus orientation by taking the mode of a circular Gaussian function fit to the neural responses. We estimated each unit’s orientation selectivity (OSI) using the following equation ^66^:

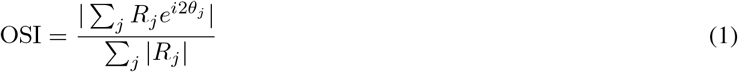

Where *R*_*j*_ is the mean firing rate and *θ*_*j*_ the orientation for the *j*^th^ stimulus. The OSI values shown in Fig. 1D were directly calculated from the observed responses. We next studied the impact of our stimulus manipulations on the units’ orientation tuning. For this analysis, we fit four circular Gaussian functions (one per stimulus family) to the responses of each unit. The Gaussian quartet shared the same mode across stimulus families, but could vary in their amplitude and bandwidth. The changes in response amplitude and selectivity reported in the results section were calculated from the fitted functions. Changes in gain variability with stimulus manipulations were computed by using gain variability estimates from fitting the modulated Poisson model ^36^ per unit and stimulus family. Finally, we asked how our stimulus manipulations impacted statistical response dependencies among pairs of neurons. For each pair of simultaneously recorded neurons, we estimated their ‘noise correlation’ by computing the Pearson correlation between their responses after removing the effects of stimulus condition on response mean and standard deviation.

### Data Analysis: population Fisher Information

For each recording, we computed population Fisher Information (FI) per stimulus family using the method proposed in ref. ^67^ (example shown in Fig. 1G). This method requires that the number of repeated trials, *T*, exceeds the number of units, *N*, by about a factor of 2: *T* > (*N* + 2)/2. For some of our populations, this requirement is not met. To circumvent this violation, we combined stimulus conditions whose orientation differed by 180° since these only differed in their drift direction. For each population, we summarized the FI per stimulus family by computing the median across stimulus orientations. Finally, to assess the impact of noise correlations, we compared FI with shuffled FI, calculated using the method proposed by ^67^.

### Stimulus encoding model

We fit responses of individual V1 units with the ‘Stochastic Normalization Model’, a model inspired by the original work of Hubel and Wiesel ^26^ and composed of elements introduced by many later studies ^20,32,47,58^. The model consists of a canonical set of linear-nonlinear operations and describes how band-pass filtered noise stimuli are transformed into the firing rate of a V1 cell. Stimuli are first processed by a linear filter whose output is half wave rectified. Following earlier modeling work that involved similar stimuli ^32^, the spatial profile of the linear filter is given by a derivative of a 2D Gaussian function. At the preferred spatial frequency, the orientation selectivity of this filter depends on the aspect ratio of the Gaussian, *α*, the order of the derivative, b, and the directional selectivity, *d*:

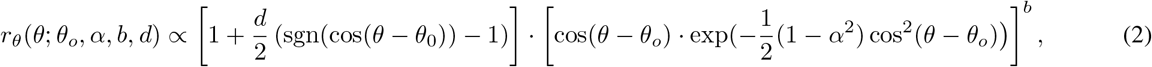

where *θ* is stimulus orientation, *θ*_*o*_ is the filter’s preferred orientation and parameter *d* ∈ [0, 1] determines direction selectivity. The function *sgn(*·) computes the sign of the argument. Because spatial frequency was not systematically varied in our stimulus set, it is not possible to uniquely determine both *α* and b from the neural responses we observed ^32^. As such, we set the derivative order to 2 unless the best fitting aspect ratio reached an upper limit of 5 – more extreme values correspond to spatial receptive fields that are atypically elongated for V1^32^. The filter’s stimulus response, *f*(*S*), was computed as the dot-product of the filter and stimulus profile in the orientation domain:

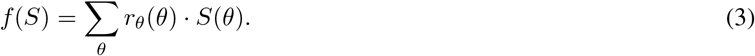

The amplitude and width of the stimulus profile directly reflect the stimulus’ contrast and dispersion, respectively. In the second stage of the model, the filter’s stimulus response is converted into a deterministic firing rate, *μ*(*S*), by subjecting the filter output to divisive normalization and passing the resulting signal through a power-law nonlinearity. This step also involves the inclusion of two sources of spontaneous discharge (one simply adds to the stimulus drive, the other is suppressed by stimuli that fail to excite the neuron) and a scaling operation:

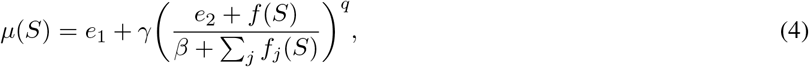

where *e*_1_ and *e*_2_ control the spontaneous discharge, *γ* the response amplitude, *q* the transduction nonlinearity, while stimulus independent constant, *β*, and the aggregated stimulus-drive of a pool of neighboring neurons, ∑_*j*_*f*_*j*_(*S*), provide the normalization signal.

Neural responses vary across repeated stimulus presentations. To capture this aspect of the data, the model describes spikes as arising from a doubly stochastic process – specifically, a Poisson process subject to “gain modulations” originating from noisy normalization signals with standard deviation *σ*_∈_ ^20,36^. Under these assumptions, spike count variance, 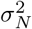, is given by:

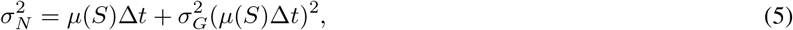

where Δt is the size of the counting window and *σ*_*G*_ the standard deviation of the response gain, given by:

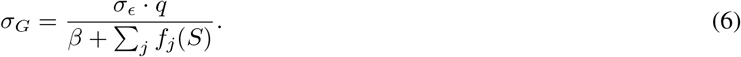

This model has 11 free parameters in total: four filter parameters (orientation preference *θ*_*o*_, spatial aspect ratio *α*, derivative order *b*, and directional selectivity *d*), four parameters controlling response range and amplitude (constant *β*, scalar *γ*, and maintained discharge *e*_1_ and *e*_2_), one parameter for the nonlinearity (exponent *q*), one parameter for the normalization noise (*σ*_∈_), and one final parameter that controlled the degree to which the normalization signal depended on stimulus dispersion. We computed the model prediction for every trial and used a Bayesian optimization algorithm to find the best fitting parameters ^68^. An example model fit is shown in Fig. 2A. To assess the goodness-of-fit, we computed the Pearson correlation between predicted and observed response mean and variance across all stimulus conditions (Fig. 2B). Units were excluded from further analysis if the Pearson correlation fell below 0.5 for response mean or below 0.2 for response variance. 352 out of 378 candidate units (93.1%) met this threshold.

### Decoding V1 population activity

We leveraged the stimulus encoding model to decode V1 population activity on a trial-by-trial basis, building on the method proposed in ref. ^20^. Specifically, assuming that the recorded neurons fire independently from one another, we modeled a pattern of spike counts {*K*_*i*_} realized during a window of length Δ*t* using a negative binomial distribution ^36^:

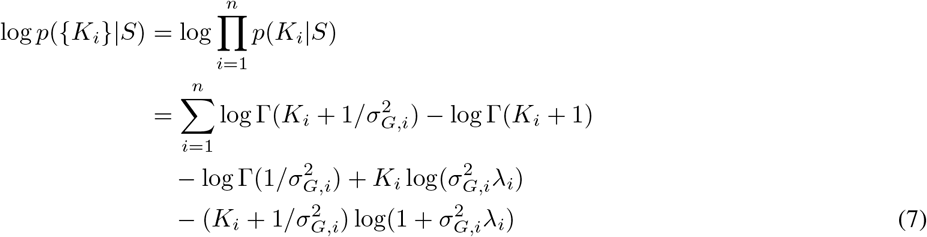

where the rate *λ*_*i*_ = *μ*_*i*_(*S*)Δ*t* and gain variability *σ*_*G,i*_ are given by the stochastic normalization model (Eq. 4 & 6).

In our experiment, the stimulus was fully defined by three parameters: its peak orientation *θ*_*S*_, its spatial contrast *c*_*S*_ and its orientation dispersion *σ*_*S*_. To obtain the orientation likelihood function for a given trial, we first calculated the likelihood of each possible parameter combination in a finely sampled 3D grid (orientation: [0:.5:360°], contrast: [0.07:.05:1.4], and dispersion: [1:3.4:99°]). We then marginalized across the contrast and dispersion dimension, yielding the orientation likelihood function (examples shown in Fig. 2D). This function typically appeared Gaussian in shape (the Pearson correlation coefficient between the likelihood function and best-fitting Gaussian was on average r = 0.985). We therefore used the peak of the best-fitting Gaussian as the point-estimate of stimulus orientation. The width of the Gaussian defines the uncertainty of this estimate. Alternative ways to quantify both statistics which did not involve fitting a Gaussian function yielded highly similar values.

### Candidate metrics of uncertainty

We evaluated different aspects of V1 population activity as proxies for the decoder’s orientation uncertainty. Specifically, we considered response magnitude, *R*_*M*_ (which indicates certainty, the inverse of uncertainty), response dispersion, *R*_*D*_, and cross-neural gain variability, *R*_*G*_, computed as:

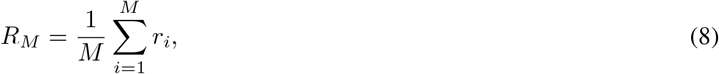

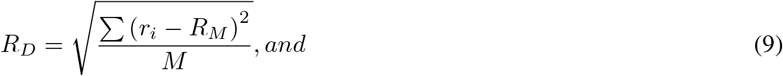

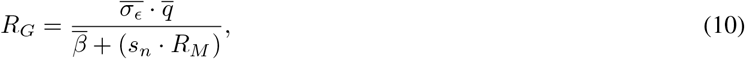

where *M* is population size, 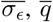, and 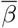 are the mean values of the corresponding parameter estimates across all units (Eq. 4 and 6), and *s*_*n*_ is a fixed scalar parameter used to relate the observed response magnitude to the unobserved stimulus-driven component of the normalization signal (its value was estimated through simulation of the stochastic normalization model).

These metrics are intended to reflect properties of the sensory population activity that informs perceptual interpretations. Due to the limited size of our recorded populations, estimated values will typically deviate from the ground truth. We therefore also considered ‘inferred estimates’ obtained by leveraging knowledge of each unit’s tuning properties. Specifically, for each trial, we estimated the most likely stimulus orientation, contrast, and dispersion under our encoding model:

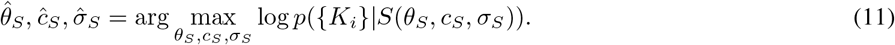

We found this maximum likelihood stimulus estimate via gradient descent using a Bayesian optimization algorithm ^68^. We then used the encoding model to compute the expected response magnitude, response dispersion, and gain variability for this specific stimulus. The first and last estimate were obtained by computing the cross-unit average of Eqs. 4 and 6, while the expected standard deviation of the population response is given by:

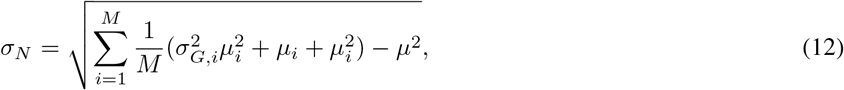

where *σ*_*G,i*_ and *μ*_*i*_ are the expected gain-variability and response average of neuron *i* and *μ* is the expected magnitude of the population response.

We examined our ability to recover each of the direct and inferred metrics from populations of various sizes simulated from the stochastic normalization model. To simulate an idealized population, for most parameters we used the median value from the model fits to our recorded units. We constrained *θ*_*o*_ to evenly tile the orientation domain and we randomly drew *γ* for each simulated unit from an exponential distribution ^53^. We simulated responses for our experiment 1 with 2 levels of contrast, 2 levels of orientation dispersion, 16 orientations, and 35 repeated trials per stimulus condition. Each direct and inferred metric was computed from these population responses as in our experimental data and then compared to the ground truth values from the simulations (results summarized in Fig. 3A).

### Estimating partial correlations

We developed a method to estimate the association between each candidate metric and orientation uncertainty while controlling for one other candidate metric (i.e., a statistic known as a ‘partial correlation’). First, we Z-scored each metric and then rank-ordered the trials as a function of the metric we sought to control for. We considered non-overlapping bins of 50 consecutive trials and computed the variance of the “frozen” metric. If this value did not exceed a threshold level of *σ*^2^ = 0.005, we proceeded to calculate the Spearman correlation between the “test” metric and the likelihood width for that set of trials. This analysis yielded a distribution of partial correlation values, shown in Fig. 4. We computed the average of each distribution on the Fisher Z-transformed values, and then used the inverse transformation to map this back onto a scale from [–1, 1] (triangles in Fig. 4B,C).

### Artificial Neural network

We trained a family of feed-forward multi-layer perceptron neural networks on trial-by-trial spike count vectors to predict trial-by-trial ground truth likelihood width estimates. A unit’s spike count was included only if a unit was present for the entire duration of the experiment. On average 78% percent of units’ spike counts were used. We implemented networks with the TensorFlow framework with the AdamW optimiser with an objective to minimize the mean squared error between ground truth and network-predicted likelihood. Training proceeded in two phases.

#### Phase 1

For each dataset, we trained 360 unique models which varied in the number of hidden layers (1, 2 or 3), number of hidden units per layer (10, 20, 30, 40, or 50), dropout rate between layers (0.05, 0.1, 0.15, or 0.2), weight decay (0.0, 0.001, or 0.001), and the learning rate (0.001 or 0.0001). For each configuration of hyper-parameters we trained five networks on 80% of trials (training/validation set) and obtained a cross-validated prediction on the held out 20% of trials, rotating trials between training and held out set such that each trial had a cross-validated prediction. With this grid search we determined the set of hyper-parameters which minimized each dataset’s held-out loss.

#### Phase 2

Optimal-hyper parameters in phase 1 differed across datasets. We used this range of optimal hyper-parameters to train 96 networks on each dataset (each one using a different hyper-parameter configuration; hidden layers: 2 or 3; hidden units: 20, 30, 40, or 50; dropout rate: 0.05, 0.1, 0.15, or 0.2; weight decay: 0.0, 0.001, or 0.001; and learning rate: 0.001) and computed their held-out loss as before. These held-out losses represent an ensemble-based estimate ^69^ of the MLP model-class’s predictive accuracy, which we report in our results.

## Acknowledgments

This work was supported by the US National Science Foundation (Graduate Research Fellowship to Z.M.B.-S.), the US National Institutes of Health (grant nos. T32 EY021462 and K99 EY032102 to C.M.Z., and EY032999 to R.L.T.G.), and the Whitehall Foundation (grant no. UTA19-000535 to R.L.T.G.). The funders had no role in the study design, data collection and analysis, decision to publish or preparation of the manuscript.

## Author Contributions

R.L.T.G, Z.M.B.-S, C.M.Z, and O.J.H participated in the conceptualization of the study. R.L.T.G supervised all aspects of the study. Z.M.B.-S. collected and analyzed experimental data. Z.M.B.-S and C.M.Z participated in the development of and analysis of computational model simulations. Z.M.B.-S and O.J.H participated in the development of and analysis of ANNs. R.L.T.G and Z.M.B.-S wrote the original manuscript draft. R.L.T.G, Z.M.B.-S, C.M.Z, and O.J.H participated in review and editing of the manuscript.

## Supplementary Information

### Further analysis of the impact of noise correlations on orientation coding

In visual cortex, trial-to-trial response fluctuations are often correlated among neurons ^39^. It is well-known that these so-called ‘noise correlations’ can impact the coding capacity of neural populations, though this need not be the case ^40^. In our data, we found little evidence for a systematic impact of noise correlations on orientation coding. The values we observed are in line with earlier findings: small, but positive on average, with a substantial spread across neuronal pairs (median values ranged from 0.11 to 0.13 across stimulus families; Supplementary Fig.1a). To assess the impact of noise correlations, we first computed Fisher Information on responses that were randomly shuffled across repeated trials of the same condition. This procedure removes noise correlations but did not systematically alter population Fisher information. This can be seen for an example population (Supplementary Fig.1b), and also held true across populations (Supplementary Fig.1c). To further test if noise correlations have an impact on coding quality, we trained ANNs to estimate stimulus orientation from single trial population responses (see Supplementary Methods). One network variant was trained on condition shuffled spike counts (“correlation blind decoder”) while the other was trained on observed spike counts (“correlation aware decoder”). Each variant was tested on unshuffled held out responses. If noise correlations impact coding capacity, we would expect the correlation blind decoder to perform systematically worse. This was not the case. The median estimation error was comparable across both decoders (Supplementary Fig.1d).

**Supplementary Figure 1.**
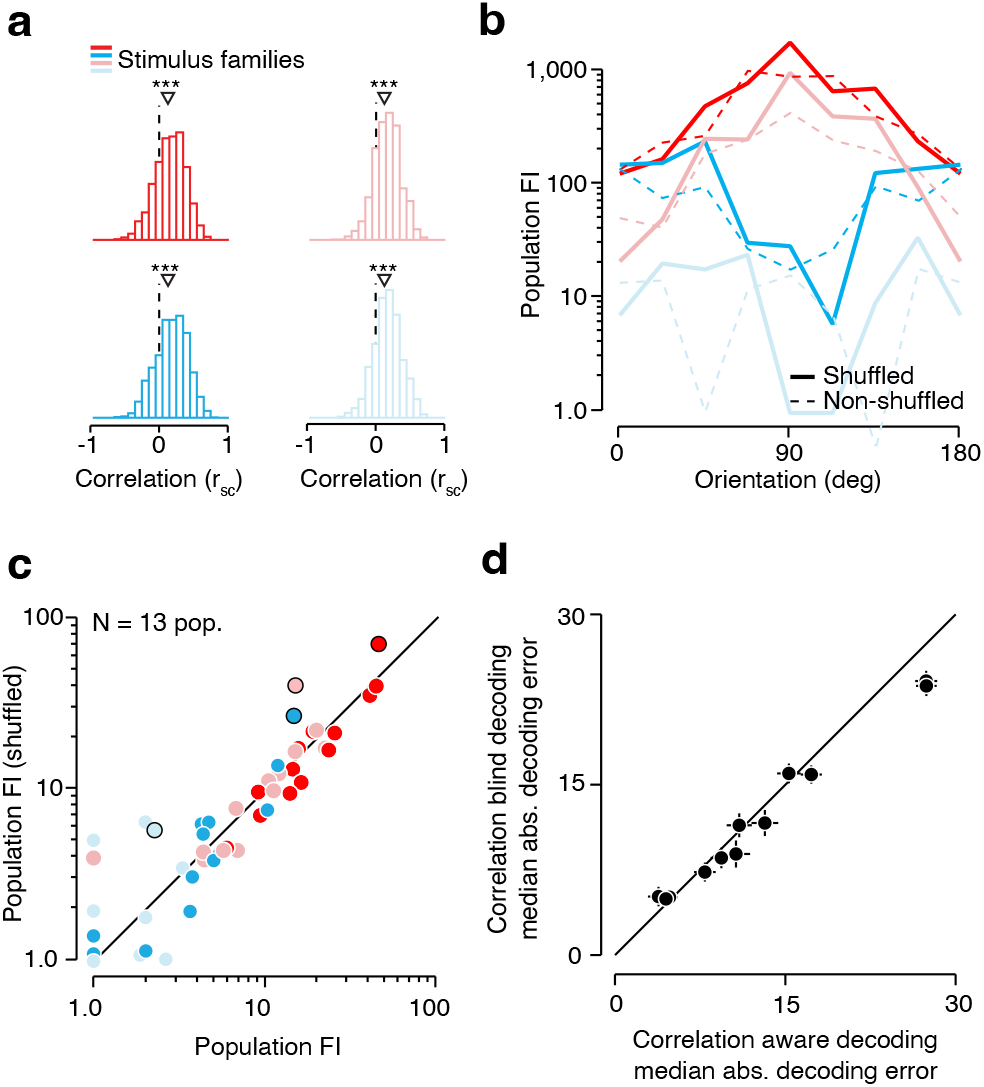
Noise correlations have negligible effect on Fisher information and estimation error (**a**) Histogram of pairwise noise correlation values for an example population (same population as shown in Fig.1). High contrast/ low orientation dispersion stimuli: median value = 0.11 (upper left, dark red); high contrast/ high dispersion stimuli: median value = 0.13 (lower left, dark blue); low contrast/ low dispersion stimuli: median = 0.12 (upper right, light red); low contrast / high dispersion stimuli: median = 0.13 (lower right, light blue). *** P < 0.001, Wilcoxon signed-rank test. (**b**) Shuffled population information (solid lines) as a function of stimulus orientation for example population. Dashed lines indicate non-shuffled population FI. (**c**) Shuffled population FI per stimulus family and population compared to population FI. (**d**) Median absolute decoding error for a correlation aware (abscissa) and correlation blind (ordinate). Each point summarizes results for a single population. One outlier population is not shown. Error bars indicate *±* SEM.

### The stochastic normalization model: parameter estimates

We fit responses of each unit with a previously proposed model of V1 function (the stochastic normalization model). This model has 11 free parameters in total (see Methods). Parameter estimates varied across units, reflecting inter-neuronal differences in the operations that shape neural stimulus selectivity, response nonlinearities, and response variability (Supplementary Fig.2).

**Supplementary Figure 2.**
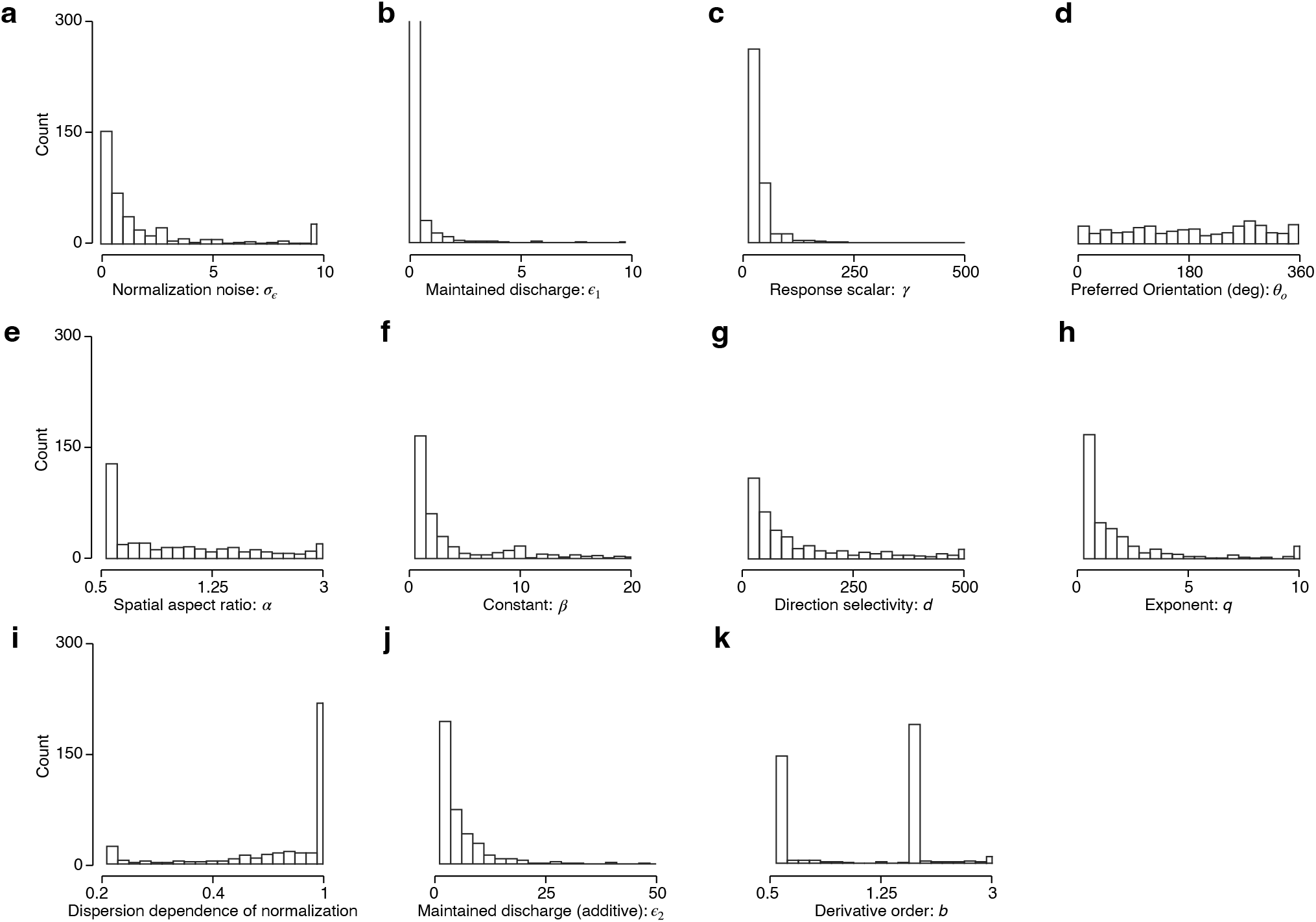
Histogram of stochastic normalization model parameters; n = 352 units. (**a**) *σ*_∈_: Normalization noise parameter. (**b**) *e*_1_: Multiplicative parameter contributing to maintained discharge. (**c**) *γ* : response scalar. (**d**) *θ*_o_: preferred orientation. (**e**) *α:* spatial aspect ratio. (**f**) *β:* stimulus independent constant. (**g**) *d*: direction selectivity. (**h**) *q* exponent. (**i**) Degree to which normalization signal depends on stimulus dispersion. Values of one indicate no effect. (**j**) *e*_2_: additive maintained discharge. (**k**) *b*: derivative order.

### The likelihood function yields unbiased stimulus orientation estimates

We developed a decoding method to infer stimulus orientation from V1 population activity on a trial-by-trial basis. For each trial, we computed the orientation likelihood function. The maximum of this function provides a point estimate for stimulus orientation, while the spread of this function indicates the uncertainty of this estimate. This decoding method is unbiased: it yields orientation estimates that, on average, are correct (Supplementary Fig.3).

**Supplementary Figure 3.**
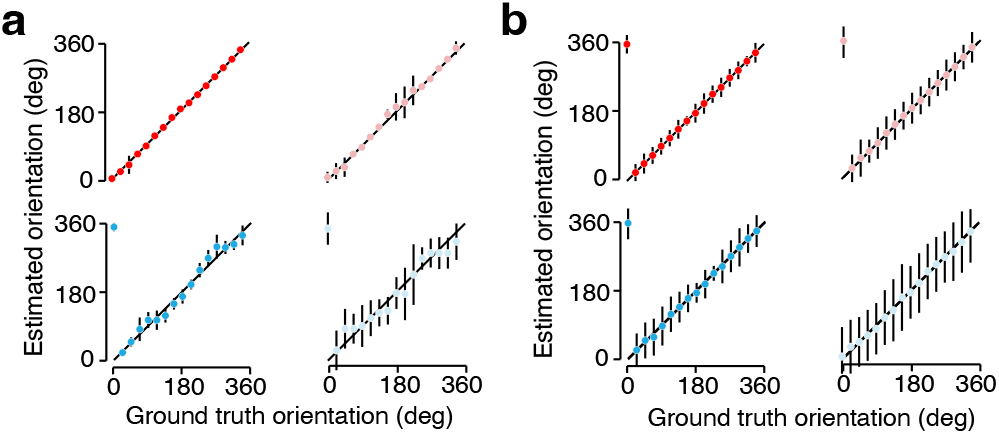
Estimating stimulus orientation from decoded V1 population activity. (**a**) Average orientation estimates as a function of ground truth stimulus orientation for an example population. Each panel summarizes results for a different stimulus family: high contrast/ low orientation dispersion stimuli (upper left, dark red); high contrast/ high dispersion stimuli (lower left, dark blue); low contrast/ low dispersion stimuli (upper right, light red); low contrast / high dispersion stimuli (lower right, light blue). Error bars indicate mean ± 1 SD. On average, orientation estimates are unbiased; all points are close to the line of unity. Note that the variance of the estimates increases with decreasing population FI. (**b**) Average orientation estimates as a function of ground truth stimulus orientation for all recordings. Error bars indicate mean estimate *±* 1 SD.

### Decoding V1 population activity under a Poisson assumption

The decoding method used in the main paper treats spikes as if they arise from a doubly stochastic process (specifically, a modulated Poisson process). We wondered how this aspect of the method influenced our findings. To address this, we decoded neural activity using a simplified variant of the method that assumes a single source of response variability (the Poisson process, see Supplementary methods). We computed the corresponding orientation likelihood function and performed the same analyses as in the main paper. Qualitatively, this procedure yielded highly similar results (Supplementary Fig.4a). Quantitatively, there was a systematic difference. The association between the candidate metrics and the width of the likelihood function was higher for the more complex decoder used in the main paper (Supplementary Fig.4b).

**Supplementary Figure 4.**
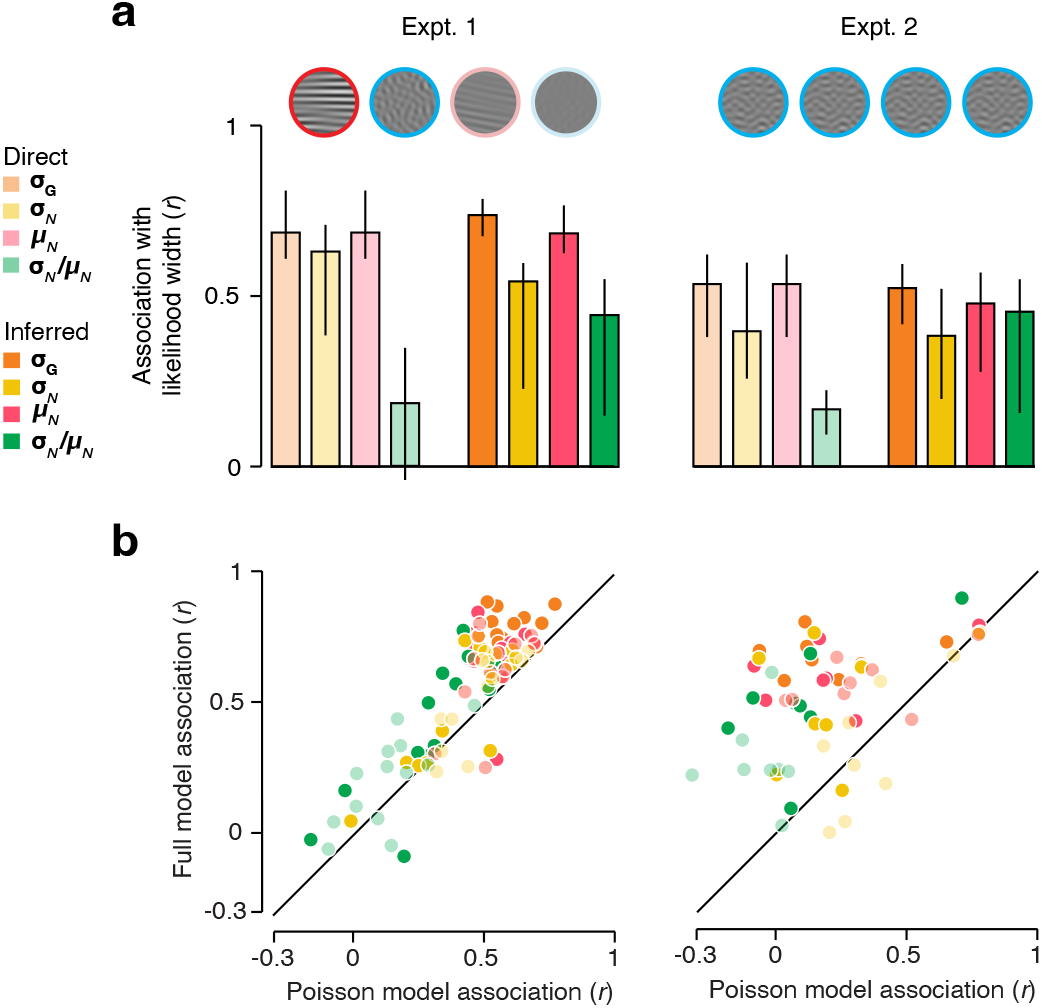
Comparison of two decoding methods. (**a**) Association between candidate uncertainty metrics and likelihood width of a Poisson decoder for experiment 1 (left) and experiment 2 (right). Error bars indicate interquartile range. (**b**) Comparison of likelihood width association for the more complex decoding method (ordinate) and the simpler Poisson variant (abscissa) for experiment 1 (left) and experiment 2 (right). Each dot represents a single candidate metric for one population.

### Further analysis of the role of inter-metric dependencies

We identified various candidate neural metrics of uncertainty that exhibited modest-to-high associations with likelihood width. These metrics were highly correlated amongst themselves. For example, in population 1, the rank correlation between inferred response magnitude and gain variability across all trials is 0.79 (Supplementary Fig.5A, left). Of course, a randomly selected subset of trials may exhibit a different level of correlation (Supplementary Fig.5A, middle). We reasoned that this sampling variability offered an opportunity to test which metric was the strongest driver of the relationship with likelihood width. Specifically, if a candidate metric is directly associated with likelihood width, the strength of this association should not systematically depend on the level of inter-metric correlation. To test this we drew 10,000 samples of 100 trials without replacement from the set of all trials for a population. Samples were included only if the difference in the variance of each metric fell below a threshold (see Supplementary Methods). We calculated the rank correlation between candidate metrics and likelihood width for each sample and sorted these values by inter-metric correlation (Supplementary Fig.5A, right). Inferred gain-variability did not depend much on inter-metric correlation, as evidence by the nearly horizontal slope of the orange line. In contrast, inferred response magnitude was only strongly associated with likelihood width when it was strongly correlated with gain variability (pink line). These results held true across all populations (Supplementary Fig.5B, left and middle), and were also evident for experiment 2 in which variability is largely due to internal fluctuations (Supplementary Fig.5C, left and middle).

**Supplementary Figure 5.**
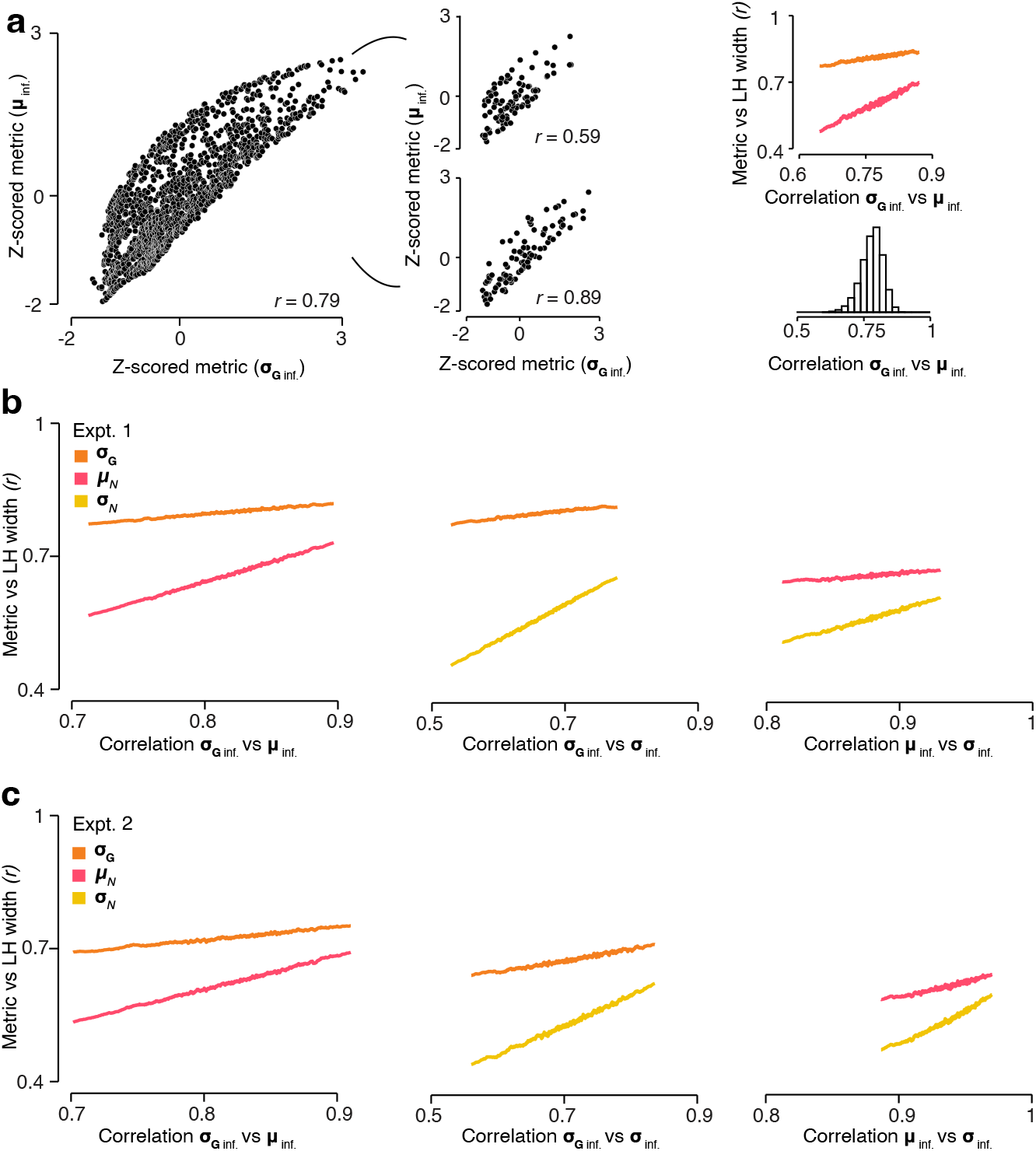
Studying inter-metric dependencies (**a**)(Left) Comparison of two z-scored inferred metrics for one example population; Ordinate: (*σ*_*Ginf*._; Abscissa: *μ*_*inf*._. Each dot is a single trial. (Middle) Two random samples of 100 trials from the example population. The variance of inferred gain variability and inferred response magnitude is 1.11 and 1.10 respectively (top) and 0.61 and 0.62 (bottom). (Right, top) Correlation of metric with likelihood width as a function of inter-metric correlation for the example population. (Right, bottom) Distribution of correlation coefficients between two candidate metrics across all 10,000 samples. (**b**) Experiment 1: Cross population analysis for *σ*_*Ginf*._ vs *μ*_*inf*._ (left), *σ*_*Ginf*._ vs *σ*_*inf*._ (middle), and *μ*_*inf*._ vs *σ*_*inf*._ (right). (**c**) Same as for **b** for experiment 2.

## SUPPLEMENTARY METHODS

### Comparing correlation-aware and correlation-blind decoders

We trained two types of feed-forward multi-layer perceptron neural networks on trial-by-trial spike count vectors to predict ground truth stimulus orientation. All networks consisted of 40 hidden units per layer, the drop out rate between layers was 0.2, the weight decay was 0.0, and the learning rate was 0.001. Input to orientation decoding networks were either spike counts which were shuffled randomly per neuron within a stimulus condition (defined as a specific combination of orientation, contrast, and dispersion) or non-shuffled responses. We then assessed the orientation estimation quality for networks trained on shuffled responses (“correlation-blind” decoders) or non-shuffled responses (“correlation-aware” decoders) on the same held out non-shuffled validation data. We implemented networks with the TensorFlow framework with the AdamW optimiser with an objective to minimize the mean squared error between ground truth and network-predicted orientation. As with the likelihood width predicting ANNs, for each configuration of hyper-parameters we trained five networks on 80% of trials (training/validation set) and obtained a cross-validated prediction on the held out 20% of trials, rotating trials between training and held out set such that each trial had a cross-validated prediction. We then computed and report held-out loss.

### Decoding V1 activity under a Poisson assumption

We used the same decoding procedure and method of calculating likelihood width as in the main paper. The key difference was that the gain-variability term in eq. 7 was set to zero.

### Analysis of role of inter-metric dependencies

In this analysis, we took a random sample of 100 trials from all trials in an experiment without replacement. We calculated the variance of two metrics for this sample of trials. If the difference in the variance of each metric was below a threshold 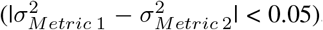, the Spearman rank correlation between each metric and the correlation of each metric with likelihood width was computed. We repeated the procedure 10,000 times.

